# The intracellular and plasma membrane pools of PI4P control megakaryocyte maturation and proplatelet formation

**DOI:** 10.1101/2022.07.28.501851

**Authors:** Ana Bura, Antonija Jurak Begonja

**Author notes:** **Correspondence:** Antonija Jurak Begonja, PhD, University of Rijeka, Department of Biotechnology, Laboratory of hematopoiesis, R. Matejcic 2, 51 000 Rijeka, Croatia.

## Abstract

Megakaryocytes (MKs) develop from hematopoietic stem cells (HSCs) after stimulation by the cytokine thrombopoietin (TPO). During megakaryopoiesis, MKs enlarge, undergo the process of endomitosis and develop intracellular membranes (the demarcation membrane system, DMS) which serve as a source for future platelets (PLTs). During DMS formation, there is an active transport from the Golgi apparatus to the DMS for the delivery of proteins, lipids, and membranes. The most important phosphoinositide that controls anterograde transport from the Golgi apparatus to the PM is phosphatidylinositol-4-monophosphate (PI4P) controlled by the SACM1L phosphatase at the Golgi and the endoplasmic reticulum (ER). The role of SACM1L and PI4P in megakaryopoiesis has not been investigated so far. Here we show that in primary mouse MKs, SACM1L is mostly localized and condensed perinuclearly in immature MKs, while at later stages it is mostly dispersed and confines to the ER. At the same time, PI4P is mostly found at the Golgi apparatus and the plasma membrane (PM) in immature MKs while in mature MKs it is in the periphery of the cell and at the PM. The exogenous expression of wild-type, but not C389S mutant (catalytically dead) SACM1L, results in the retention of the Golgi apparatus leading to the increased number of immature MKs, as well as a decreased number of MKs that form proplatelets. The inhibition of the production of PI4P specifically at the PM (inhibiting PI4-kinase IIIα) resulted also in a significant decrease of MKs that form proplatelets. These results indicate that both Golgi and PM pools of PI4P mediate MK maturation and proplatelet formation.

## Introduction

Megakaryocytes (MKs) are the largest (50-100 μm in diameter) and one of the rarest (0.01%) cells in the bone marrow (BM)^1^. They differentiate from hematopoietic stem cells (HSCs) in response to thrombopoietin (TPO), a major cytokine that is produced by the liver and binds to its receptor c-Mpl promoting the development and growth of MKs^1,2^. During megakaryopoiesis, MKs undergo the process of endomitosis, increase in size, pack proteins into α- and dense granules, and develop a system of membranes called the demarcation membrane system (DMS) that is a source of membranes for future platelets (PLTs)^3^. The DMS is composed of numerous cisternae and tubules that are continuous with the PM and it requires high levels of proteins and lipids for its formation^1^. The formation of DMS starts in the perinuclear region, near the Golgi apparatus^4^. It has been shown that the vesicles from the *trans*-Golgi network (TGN) are localized close to the DMS, and fuse with the DMS, while the blockage of Golgi trafficking increases the number of immature MKs^4^. In the later stages of DMS formation, the endoplasmic reticulum (ER) makes membrane contact sites (MCS) with the DMS^4^. This suggests that vesicles for cargo transport deliver necessary proteins, lipids, and membranes for the formation of DMS. One of the most abundant phosphoinositides (PIs) at the Golgi apparatus and the ER is phosphatidylinositol-4-monophosphate (PI4P)^5^. By binding effector proteins, PI4P regulates the anterograde transport of vesicles from the TGN to the plasma membrane (PM)^6–8^. PI4P recruits clathrin adaptors and Golgi-localized Gamma-adaptin ear homology, Arf-binding proteins (GGAs)^9^.

SACM1L (suppressor of actin mutations 1-like protein) is a SAC1 domain-containing phosphatase^10^ that mainly dephosphorylates PI4P at the Golgi apparatus and the ER^5^. It is a transmembrane protein with a CX5R(T/S) catalytic motif^11^ that localizes to both ER and Golgi which is growth factor-dependent^12^. When the cells are stimulated with growth factors, SACM1L is mostly found at the ER resulting in the accumulation of PI4P at the

Golgi apparatus and stimulation of the anterograde transport from the Golgi to the PM. In growth factor deprived conditions, SACM1L mostly shifts from the ER to the Golgi apparatus where it dephosphorylates PI4P resulting in a decrease of the anterograde trafficking from the Golgi apparatus to the PM.

Data from the human and mouse PLT proteome revealed that SACM1L is highly expressed in human (5800 copies) and mouse (10500 copies) PLTs^13^ suggesting high copy numbers in MKs as well. Considering the proximity of the Golgi apparatus to maturing DMS and the contribution of its transport vesicles that supply growing membranes, and the PI4P that regulates anterograde Golgi trafficking, we studied the role of PI4P and SACM1L in primary mouse MKs and DAMI cell line.

Here we show that in primary mouse MKs, PI4P translocates from the Golgi apparatus to the PM during MK maturation and that the exogenous expression of wild-type SACM1L results in smaller MKs that produce fewer proplatelets. Since the inhibition of PI4P production at the PM by phosphoinositol-4 kinase IIIα (PI4KIIIα) also results in a decrease in proplatelet formation we suggest that both the Golgi and PM pool of PI4P mediate MKs maturation and proplatelet formation.

## Materials and methods

### Antibodies

Primary antibodies were obtained from the following resources: polyclonal rabbit anti-SACM1L (ABIN949071) was from antibodies-online, monoclonal mouse anti-PDIA3 (AMAb90988) was from Atlas Antibodies, monoclonal mouse anti-GM130 (610823) was from BD Transduction Laboratories, monoclonal mouse anti-PI4P IgM (Z-P004) was from Echelon Biosciences, monoclonal mouse anti-KDEL (sc-58774) was from Santa Cruz Biotechnology, polyclonal rabbit anti-TGN46 (ab16059) was from Abcam, monoclonal rat anti-GPIbβ (M050-1) was from Emfret Analytics, rabbit anti-β1 tubulin was a kind gift of dr. Joe Italiano (Brigham&Women’s Hospital, Harvard Medical School, Boston), phalloidin (A12379) conjugated with Alexa Fluor (AF)-488 was from Invitrogen, monoclonal mouse anti-GAPDH (MAB374) was from EMD Millipore. Secondary antibodies were obtained from the following resources: goat anti-mouse conjugated with Alexa Fluor (AF)-555 (A21429), goat anti-rabbit conjugated with Alexa Fluor (AF)-555 (A21429), and goat anti-mouse conjugated with Alexa Fluor (AF)-488 (A11029) were from Life Technologies, goat anti-mouse conjugated with Alexa Fluor (AF)-568 (A21043), goat anti-rabbit conjugated with Alexa Fluor (AF)-488 (A11070), and goat anti-rat conjugated with Alexa Fluor (AF)-650 (SA5-10021) were from Invitrogen, goat anti-rabbit IgG HRP-linked (7074S) and goat anti-mouse IgG HRP-linked (7076S) were from Cell Signaling.

### DAMI and HEK293T cells

DAMI cells were grown in RPMI 1640 medium (Roswell Park Memorial Institute, Lonza) supplemented with 10% fetal bovine serum (FBS, Pan Biotech) and 1% penicillin/streptomycin (Lonza). On day 0, DAMI cells were differentiated with 100 nM phorbol 12-myristate 13-acetate (PMA, P8139-1MG, Sigma Aldrich) and cultivated for 4, 8, 24, and 48 hours in full medium or 4, 8, and 48 hours in medium lacking 10% FBS. For transfection, DAMI cells were seeded and differentiated with 100 nM PMA and after 24h transfected with pEGFP-SACM1L, pEGFP-C389S-SACM1L (a kind gift of dr. Gerry Hammond, Department of Cell Biology, University of Pittsburgh School of Medicine, Pittsburgh, PA, USA) or pmCherry-PI4P-SidM (a kind gift of dr. Antonella De Matteis, Telethon Institute of Genetics and Medicine Pozzuoli, Naples, Italy; Department of Molecular Medicine and Medical Biotechnology University of Napoli Federico II, Naples, Italy) with Lipofectamine 3000 transfection reagent. 24h after transfection the cells were fixed and stained for proteins of interest. HEK293T cells were grown in DMEM (Dulbecco’s modified Eagle’s medium, Pan Biotech) supplemented with 10% FBS (Pan Biotech) and 1% penicillin/streptomycin (Lonza). Cells were seeded in full medium for 24h before lysing for Western blot assay.

### Mice

C57BL/6J mice were used for all experiments. Mice were handled following the European Communities Council Directive of 24 November 1986 (86/609/EEC) and with institutional and national guidelines. All experimental procedures were approved by the Ethics Committees of the Department of Biotechnology and Medical School of the University of Rijeka (003-08/15-01/34; 322-01/22-01/02), as well as the Ministry of Agriculture of the Republic of Croatia (UP/I-322-01/15-01/123).

### Mouse BM and FL isolation and MK production

Mouse BM from 8-12 weeks old mice and FL from 13.5 dpc old embryos were used. Cell aggregates from the BM were dissociated by mechanical aspiration ten times through a 21G x 1” needle, ten times through a 23G x 1” needle and two times through a 25G x 5/8” needle. Dissociated cells from the BM and FL were filtered through a 70 μm cell strainer (Falcon) and centrifuged for 5 min at 1000 rpm. The cells were then resuspended and cultured in DMEM (Pan Biotech) supplemented with 10% FBS (Pan Biotech) and 1% penicillin/streptomycin (Lonza). For MK production, TPO (2% of the culture-producing cytokine) was added. Cells were cultivated for different periods as indicated in the figures and figure legends. MKs were enriched through bovine serum albumin (BSA) gradient on days 2 and 4 (FL-derived MKs) or days 3 and 5 (BM-derived MKs) for further analysis.

### Proplatelet formation assay

On day 4 after isolation of FL-derived MKs BSA gradient was performed and 18h later the number of MKs forming proplatelets was counted and shown as the percentage of MKs that form proplatelets over all MKs. The counting of MKs and proplatelets was performed using an epifluorescent microscope (Olympus) or a bright-field microscope (Zeiss) at 20x magnification. For counting the number of MKs that form proplatelets after the inhibition of PI4KIIIα, 100 nM GSK-A1 (SYN-1219-M005, Adipogen) was used.

### Mouse platelet isolation

Mouse platelets were isolated from mouse peripheral blood. Peripheral blood was drawn in 1/10 of Aster-Jandl anticoagulant. Blood was centrifuged for 8 min at 100g. PRP was diluted 1:5 in washing buffer (140 mM NaCl, 5 mM KCl, 12 mM trisodium citrate, 10 mM glucose, 12.5 mM sucrose, pH=6) and centrifuged for 6 min at 100g. PRP supernatant was taken and centrifuged for 5 min at 1200g. The pellet was resuspended in 1 mL of washing buffer and again centrifuged for 5 min at 1200g. The pellet was resuspended in 200 μL of resuspension buffer (10 mM HEPES, 140 mM NaCl, 3 mM KCl, 0.5 mM MgCl_2_, 0.5 mM sodium hydrogen carbonate, 10 mM glucose, pH=7.4). PLTs were then left to rest for 30 minutes at 37°C before lysing for Western blot assay.

### Constructs, retroviral production, and MK transduction

SACM1L from pEGFP-SACM1L and C389S-SACM1L from pEGFP-C389S-SACM1L was amplified by PCR with primers Fwd 5’ TAGGCGCCGGAATTAGATCACCGGTATGGTGAGCAAGGGCGAG 3’ and Rev 5’ ATCGGCCGCCAGTGTGCTGGAATTCAGTCTATCTTTTCTTTCTGGACC 3’. Both constructs were subcloned into a retroviral Murine Stem Cell Virus (MSCV) vector expressing EGFP into AgeI and EcoRI sites. For cloning of both constructs, NEBuilder HiFi DNA Assembly DNA Master Mix (E2621L, NEB) was used. To produce retroviruses, HEK293T cells were seeded for 24h and then co-transfected with MSCV constructs (EGFP-MSCV, EGFP-SACM1L-MSCV, or EGFP-C389S-SACM1L-MSCV) and pCL-Eco packaging plasmid using the transfection reagent Lipofectamine 3000. After 24h the medium was changed and after 72h the viral supernatants were collected. Viral supernatants were filtered through a 0.45 μm filter and stored at -80°C. On day 2 of FL culture, the cells were transduced by adding retroviral supernatants in the presence of 8 mg/mL polybrene (H9268, Sigma Aldrich). The samples were centrifuged at 800g for 30 min and incubated at 37°C for 3h. After replacement with fresh media, MKs were cultured for additional two days. On day 3, FL-derived MKs were separated via BSA gradient and analyzed for the ability to form proplatelets on day 4. The percentage of MKs that produce proplatelets was calculated from the number of EGFP+ MKs producing proplatelets over the number of all EGFP+ MKs.

### Immunostaining

DAMI cells were grown directly on coverslips while MKs were spun down on glass coverslips using the Hettich cytospin centrifuge. For SACM1L with Golgi or ER markers, the cells were fixed with 4% paraformaldehyde (PFA) in PBS for 15 min at RT, rinsed three times with PBS and permeabilized and blocked with 1% goat serum/0.2% BSA/0.05% saponin for 45 min. The cells were incubated with primary antibodies (diluted 1:100) overnight at 4°C or for 1h at RT depending on the manufacturer’s instructions. The cells were rinsed 3 times with PBS and incubated with secondary antibodies (diluted 1:500) for 1h at RT. The cells were rinsed again 3 times with PBS, stained with DAPI (4’,6-diamidino-2-phenylindole) in PBS for 1 min, rinsed, and mounted.

Immunostaining of intracellular pools of PI4P, as well as Golgi markers through MK development, was performed as previously described^14^. Briefly, all steps were performed at room temperature. Cells were fixed with 4% paraformaldehyde (PFA) in PBS for 15 min, rinsed with PBS containing 50 mM NH4Cl, and permeabilized with 0.01% digitonin in buffer A (20 mM Pipes, pH 6.8, 137 mM NaCl, 2.7 mM KCl) for 5 min. Cells were then rinsed with buffer A, blocked (5% goat serum in PBS with 50 mM NH_4_Cl) for 45 minutes, and labelled with primary antibodies diluted 1:100. The cells were then washed with buffer A, stained with secondary antibodies (1:500), post-fixed for 5 min in 2% PFA, and mounted.

The staining of the PM pool of PI4P was performed with the modified protocol as described previously^15^. Briefly, cells were fixed with 4% PFA in PBS with 0.2% glutaraldehyde (GA) for 15 min and rinsed with PBS containing 50 mM of NH4Cl. All subsequent steps were performed on ice with all pre-chilled solutions. Cells were blocked and permeabilized with 0.5% saponin (in buffer A containing 5% goat serum and 50 mM of NH4Cl) for 5 min and blocked with buffer A containing 5% goat serum and 50 mM of NH4Cl for 45 min. Then, cells were labelled with primary antibodies diluted 1:100 (anti-PI4P) or 1:200 (anti-GPIbβ), washed with buffer A, stained with secondary antibodies (1:500), post-fixed for 10 min in 2% PFA on ice and an additional 5 min on room temperature, and mounted.

For the staining of proplatelets, cells were fixed with 4% paraformaldehyde (PFA) in PBS for 15 min, rinsed with PBS, and permeabilized with 0.01% Triton X-100 in PBS for 10 min. Cells were then blocked (5% goat serum in PBS) for 45 minutes and labelled with primary antibodies diluted 1:100 or with phalloidin 488 and anti-GPIbβ (1:200). The cells were then washed with PBS, stained with secondary antibodies (1:500), and mounted.

### Epifluorescent and confocal microscopy

Epifluorescent images were obtained using an IX83 Olympus microscope equipped with a Hamamatsu Orca R2 CCD camera. The objective used was a PLAPON 60x/1.42 Oil (Olympus). The Pearson correlation coefficient and the area of the cell were measured in FIJI ImageJ^16^ software.

Confocal images were obtained using an LSM880 confocal microscope (Carl Zeiss) equipped with an Argon-laser multiline (458/488/514 nm), HeNe laser (543 nm and 633 nm). The objective used was a Plan-Apochromat 63×/1.40 oil DIC III (Carl Zeiss). For ultrastructure visualization, the airy scan mode was used. Total fluorescence intensities (TFI) and the area of cells were measured using ZEN Black software (Carl Zeiss). For fluorescence intensity measurements, imaging conditions between different samples were kept constant. Fluorescence intensity was always expressed over the area (size) of cells.

### Western blot

Cells were lysed with 3x SDS STOP/β-mercaptoethanol (200 mM Tris, 6% SDS, 15% glycerol, 10 mg bromophenol blue [pH 6.7]). Lysates were denatured for 5 min at 95 °C and proteins were separated with sodium dodecyl sulfate gel electrophoresis (SDS-PAGE) on 10% polyacrylamide gel at 100 V for 1 h in the Bio-Rad system for vertical electrophoresis. For protein size tracking PageRuler™ Plus Prestained Protein Ladder was used (Thermo Scientific, 26619). Proteins from the gel were electro-transferred onto the nitrocellulose membrane (Santa Cruz Biotechnology) in the Bio-Rad system at 0.45 A for 90 min. Membranes were blocked for 45 min in BSA/TBST (3% bovine serum albumin [Pan Biotech, P06-1391100]/Tris, NaCl, 0.1% Tween 20) and incubated with primary antibodies overnight at 4°C. Membranes were then washed 3x for 10 min with TBST and secondary antibodies were added. Protein bands were visualized with ECL™ Prime Western Blotting detection reagents (GE Healthcare Life Sciences, RPN2236) and the pictures were taken with Bio-Rad ChemiDoc™ MP Imaging System.

### Statistical analysis

All experiments were performed at least in triplicate, and data are represented as mean ± standard error of the mean (SEM). Data were analyzed by t-test or ANOVA using Prism software (GraphPad). Differences were considered significant when P values were < 0.05 (* p < 0.05; ** p < 0.01; *** p < 0.001; **** p < 0.0001).

## Results

### PI4P translocates from the Golgi apparatus towards the plasma membrane during MK maturation

Firstly, we investigated the expression and localization of SACM1L in primary MKs. We cultivated MKs from mice BM for three or five days to produce immature or mature MKs, respectively (described in detail in^17^). Briefly, immature MKs are significantly smaller than mature MKs and they have a small nuclear/cytoplasmic ratio, while mature MKs grow 2-4 fold, have a large nuclear/cytoplasmic ratio, and increase the expression of GPIbβ. In BM-derived MKs we could observe two SACM1L isoforms, a highly expressed 67 kDa isoform and an isoform of 55.5 kDa, with lower expression (Figure 1A). Since the isoform of 67 kDa was the one with the highest expression, we further on quantified the expression levels of only this isoform. The quantification of the 67 kDa SACM1L showed that there was no change during MK maturation (Figure 1B). The same isoforms were observed in human PLTs (except for the 55 kDa isoform) and control HEK293T cells. Since MKs were cultivated in a nutrient-rich environment, we expected SACM1L to localize at the ER which we confirmed in both immature and mature MKs (Figure 1C).

**Figure 1.**
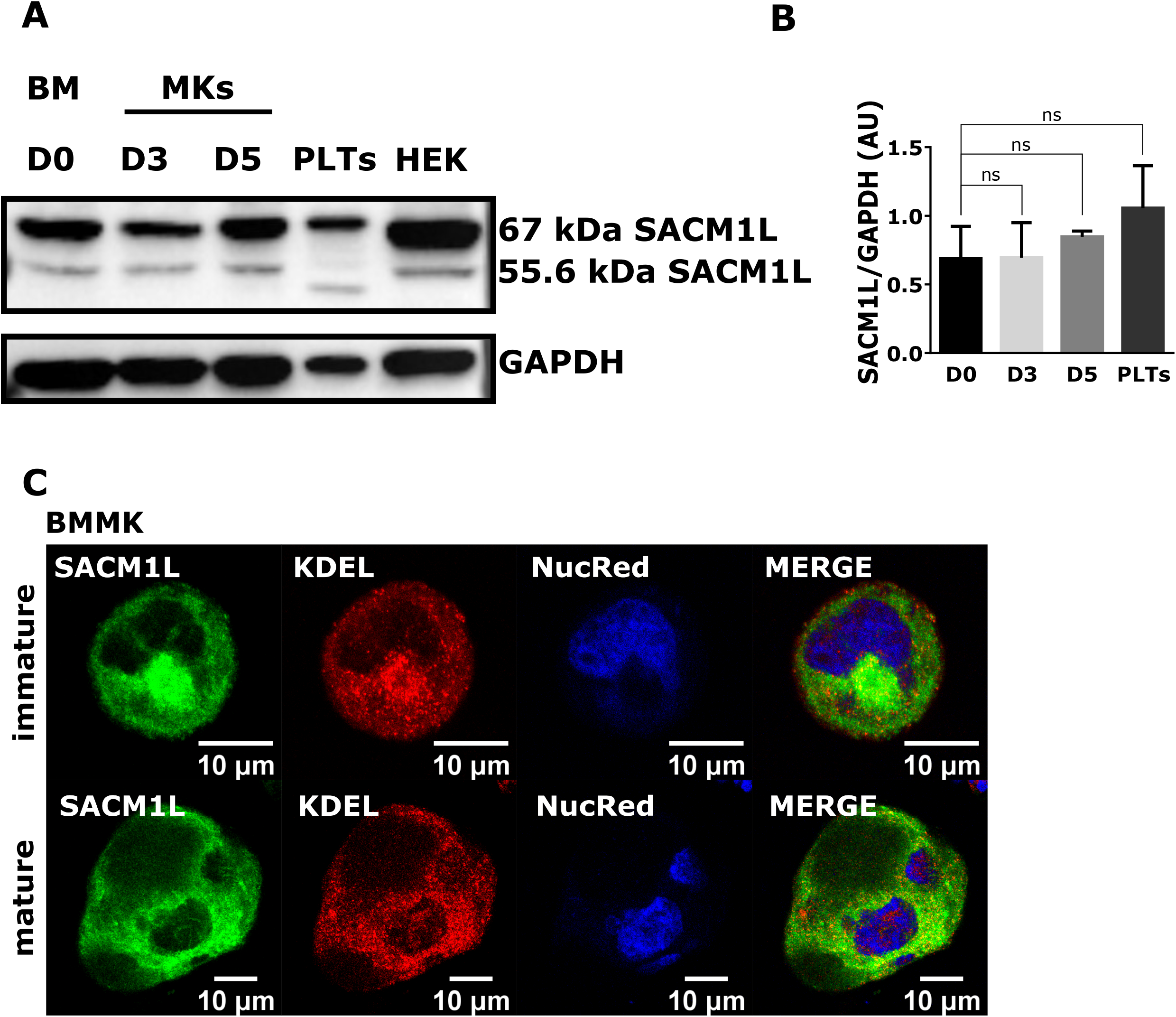
SACM1L is highly expressed in BM-derived MKs, and it is localized to the ER. MKs were isolated from the mouse BM, cultivated for 3 or 5 days, and enriched by a BSA gradient. PLTs were isolated from mouse peripheral blood, while HEK293T cells were seeded and cultured for 24h. **(A)** The cells were lysed, separated by SDS-PAGE, blotted onto a nitrocellulose membrane, and incubated with indicated antibodies. **(B)** Results in the graph are presented as means, error bars denote ± SEM from at least 3 independent experiments. *, p < 0.05; **, p < 0.01; ***, p < 0.001; **** p < 0.0001; n.s., non-significant. **(C)** MKs were fixed and stained for SACM1L and KDEL. Representative images display a single confocal optical section. The scale bar of the images is 10 μm.

In other cells, PI4P is mostly localized on the Golgi apparatus but it can also be found at the PM^5^. In immature BM-derived MKs PI4P indeed localized both at the Golgi apparatus, as shown by the colocalization with TGN46 and at the cell periphery and the PM (Figure 2A, upper panel). However, in mature MKs, it localized mostly at the periphery of the cell and the PM (Figure 2A, lower panel). Quantification of the percentage of cells that show perinuclear and PM or only PM staining of PI4P showed that approximately 90% of immature BM-derived MKs have perinuclear and PM PI4P pools, while approximately 80% of mature MKs had only PM PI4P pool (Figure 2B). This difference in PI4P localization was not specific to BM-derived MKs as it was also observed in fetal liver (FL) derived MKs cultivated for two (immature) or four days (mature)^17^. As in BM-derived MKs, in immature FL-derived MKs PI4P localized perinuclearly at the Golgi apparatus and the PM (Figure 2C, upper panel) while in mature FL-derived MKs it mostly localized at the periphery of the cell and the PM (Figure 2C, lower panel). Here the quantification revealed that approximately 80% of immature FL-derived MKs show perinuclear and PM PI4P staining while approximately 70% of mature FL-derived MKs show only PM PI4P staining (Figure 2D). To confirm that PI4P localizes at the PM, we co-stained mature MKs with GPIbβ and observed PI4P/GPIbβ colocalization at the cell periphery (Figure 2E).

**Figure 2.**
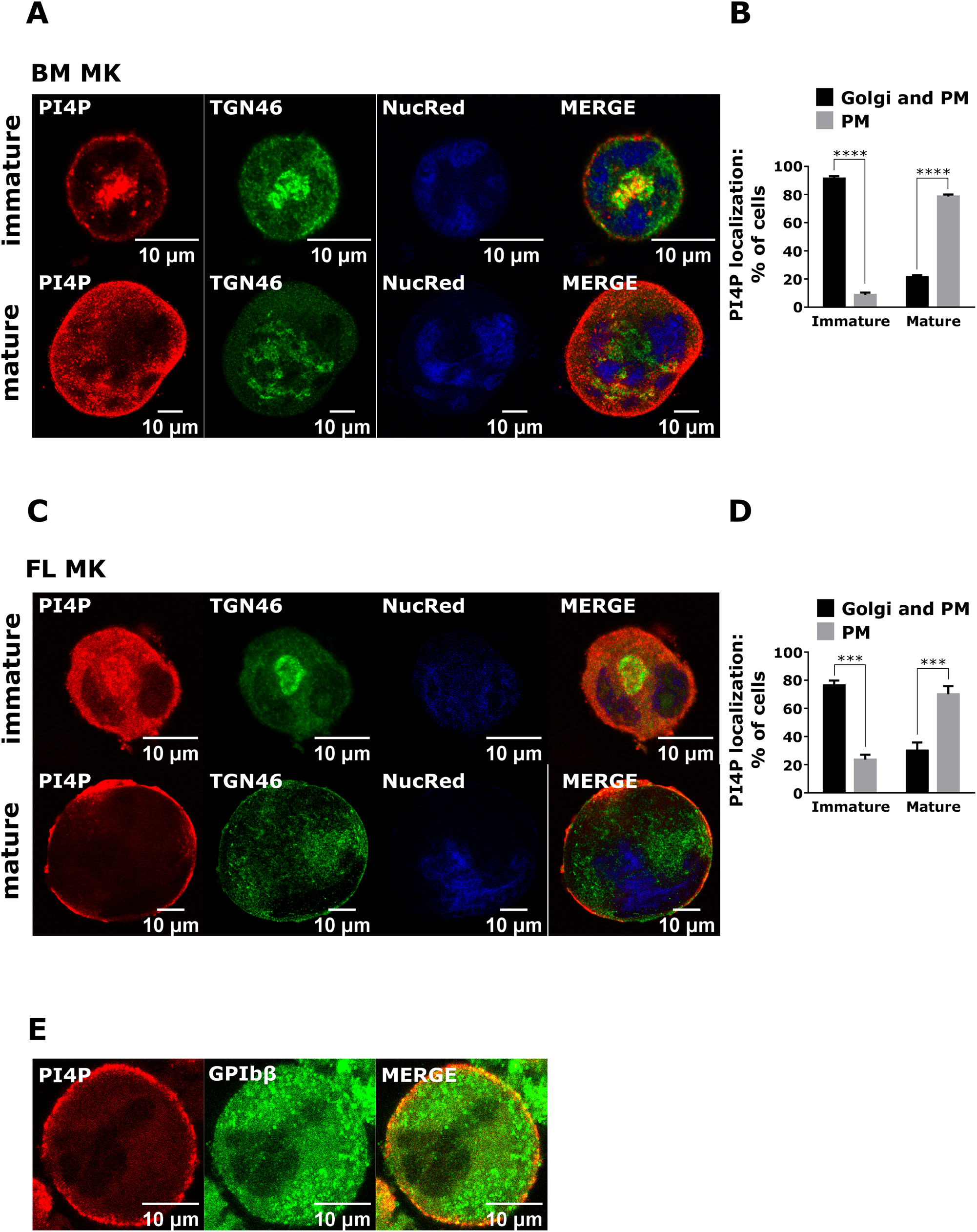
PI4P translocates from the Golgi apparatus towards the periphery of the cell and the PM in MKs. MKs were isolated from **(A)** the mouse BM or **(C** and **E)** the mouse FL, cultivated for **(A)** 3 and 5 days, **(C)** 2 and 4 days, or **(E)** four days, and enriched by a BSA gradient. The cells were then fixed, stained for PI4P and co-stained for TGN46 or GPIbβ. Representative images display a single confocal optical section. The scale bar of the images is 10 μm. **(B** and **D)** The number of MKs with different PI4P localization was counted. Results in the graphs are presented as means, error bars denote ± SEM from at least 3 independent experiments. *, p < 0.05; **, p < 0.01; ***, p < 0.001; **** p < 0.0001; n.s., non-significant.

The change in PI4P localization could be due to the changes in the Golgi morphology itself. As shown in Supplemental Figure 1, the Golgi apparatus shown by the staining for *trans*-(TGN46) and *cis*-Golgi (GM130) in immature BM- and FL-derived MKs was localized perinuclearly while in mature BM- and FL-derived MKs was dispersed in vesicular structures, as previously shown^4^. In BM-derived MKs, approximately 70% of immature MKs showed perinuclear localization of the Golgi apparatus while approximately 80% of mature MKs had dispersed Golgi into cytoplasmic vesicles (Supplemental Figure 1B). Similar changes in the Golgi morphology happen in FL-derived MKs where approximately 75% of immature MKs had perinuclear localization of the Golgi while approximately 80% of mature MKs had Golgi structures dispersed into cytoplasmic vesicles (Supplemental Figure 1D).

### The expression of wild-type SACM1L negatively impacts megakaryocyte maturation and proplatelet formation

To determine the role of the Golgi pool of PI4P that is controlled by SACM1L we expressed wild-type (WT) and catalytically inactive C389S mutant SACM1L from retroviral vector in FL-derived MKs and assessed MKs size, PI4P, and the ability of MKs to form proplatelets. MKs were readily infected with the control EGFP as well as WT or mutant SACM1L constructs with no significant difference in expression rate (Supplemental Figure 2, A and B). Both constructs were localized near the nucleus where the ER and the Golgi apparatus are expected to be (Figure 3A). Interestingly, there was no significant difference in the PI4P fluorescence intensity when WT or mutant SACM1L were expressed in MKs (Figure 3, A and B). However, MKs expressing WT SACM1L showed PI4P staining more at the perinuclear region and the PM, opposing to control MKs that showed mostly PM staining (Figure 3D). This was consistent with their maturation state since MKs expressing WT SACM1L were significantly smaller than control MKs expressing EGFP or mutant SACM1L (Figure 3C). To determine the maturation state of Golgi in MKs, we stained them with TGN46, together with MK maturation marker GPIbβ hypothesizing that if MKs are not fully mature, the Golgi apparatus would be localized more perinuclearly and less dispersed into cytoplasmic vesicles while GPIbβ would be more at the PM and less throughout the cells. MKs that were expressing WT SACM1L had a more perinuclear Golgi (∼70% of MKs), while the control MKs (90%) and the ones expressing mutant SACM1L (80%) had dispersed Golgi vesicles (Figure 3, E and F). The difference in the GPIbβ localization was less dramatic however, GPIbβ localized more at the PM in MKs expressing WT SACM1L, also suggesting that they are less mature (Figure 3, E and G). Furthermore, MK expressing WT SACML had slightly, although statistically insignificantly, increased GPIbβ TFI (Supplemental Figure 2C). This is also consistent with the hypothesis that they are smaller and less mature, as shown previously^17^.

**Figure 3.**
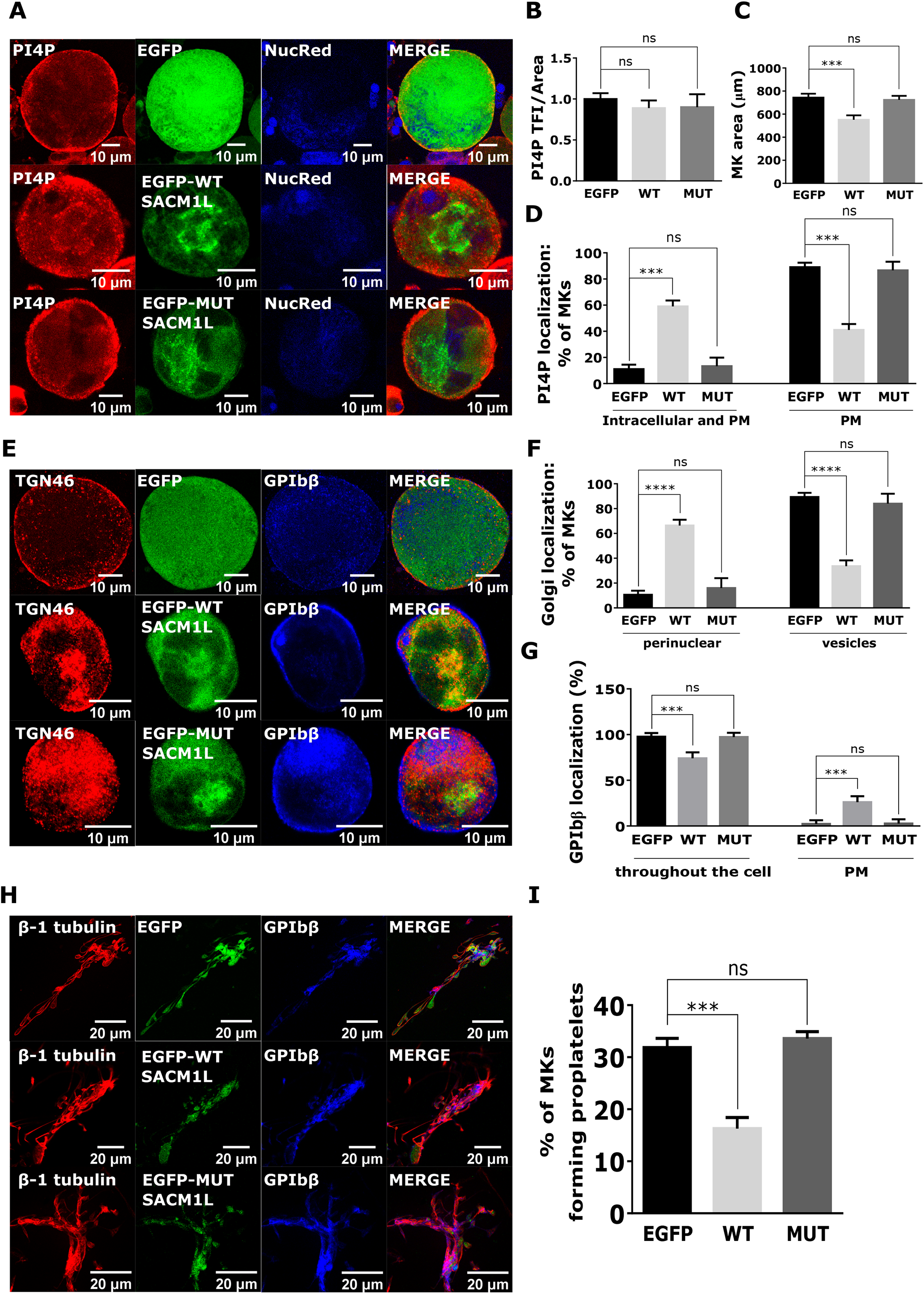
The expression of wild-type SACM1L negatively impacts MK maturation and proplatelet formation. MKs were isolated from the mouse FL. On day 2 of the culture, they were infected with wild-type or mutant SACM1L. On day 3 of the culture, they were enriched by a BSA gradient. On day 4 of the culture, they were fixed and stained for **(A)** PI4P or **(E)** TGN46 and GPIbβ. **(B)** PI4P total fluorescence intensity and **(C)** MKs area were measured using Zen black software. The number of MKs with **(D)** different PI4P localization, **(F)** different Golgi structures, **(G)** different GPIbβ localization, and **(I)** proplatelets was counted. The scale bar of the images is 10 μm. **(H)** On day 4 of the culture, proplatelets were fixed, stained for β-1 tubulin and co-stained for GPIbβ. Representative images display a single confocal optical section. The scale bar of the images is **(A** and **E)** 10 μm and **(H)** 20 μm. Results in the graphs are presented as means, error bars denote ± SEM from at least 3 independent experiments. *, p < 0.05; **, p < 0.01; ***, p < 0.001; **** p < 0.0001; n.s., non-significant.

### PI4P is necessary for proplatelet formation

We performed a proplatelet formation assay to determine the ability of SACM1L expressing MKs to produce proplatelets. MKs expressing WT SACM1L produced significantly fewer proplatelets than MKs expressing only EGFP or mutant SACM1L with no apparent change in proplatelet morphology (Figure 3, H and I). These data suggest that the Golgi pool of PI4P controlled by the SACM1L phosphatase is important for proplatelet formation and that the perturbations in PI4P levels or its localization do not allow full MK maturation and proplatelet formation.

Since PI4P localizes more on the PM with MK maturation, we wanted to see if the PM pool of PI4P has an important role in proplatelet formation. Phosphatidylinositol 4-kinase alpha (PI4KIIIα) was shown to predominantly produce PI4P at the PM in other cells^18^. Therefore, we pharmacologically inhibited PI4KIIIα with GSK-A1 and assessed PI4P intensity and the ability of FL-derived MKs to produce proplatelets. For PI4P visualization, we used two staining procedures, one that enables the imaging of predominantly the Golgi pool of PI4P and one that enables the visualization of the PM pool of PI4P^14^. In MKs that were treated overnight with GSK-A1 (100 nM), there was no difference in the fluorescence intensity of the Golgi pool of PI4P (Figure 4, A and B) with no apparent changes in the localization of TGN46 or GPIbβ. However, PI4KIIIα-inhibited MKs had significantly lower PI4P fluorescence intensity at the PM than the control MKs (Figure 4, C and D). Here, we do not show the TGN46 or GPIbβ staining since the PM staining protocol was not suitable for the detection of these proteins, which are found intracellularly. The change in PI4P levels is consistent with the expected localization of PI4KIIIα. Moreover, MKs treated with the PI4KIIIα inhibitor were smaller (Figure 4F) and produced significantly fewer proplatelets than the control MKs (Figure 4H). However, similar to the results obtained when expressing wild-type SACM1L, the morphology of the proplatelets produced by MK that were inhibited for PI4KIIIα did not change (Figure 4, E and G). These data suggest that the PM pool of PI4P also contributes to proplatelet formation.

**Figure 4.**
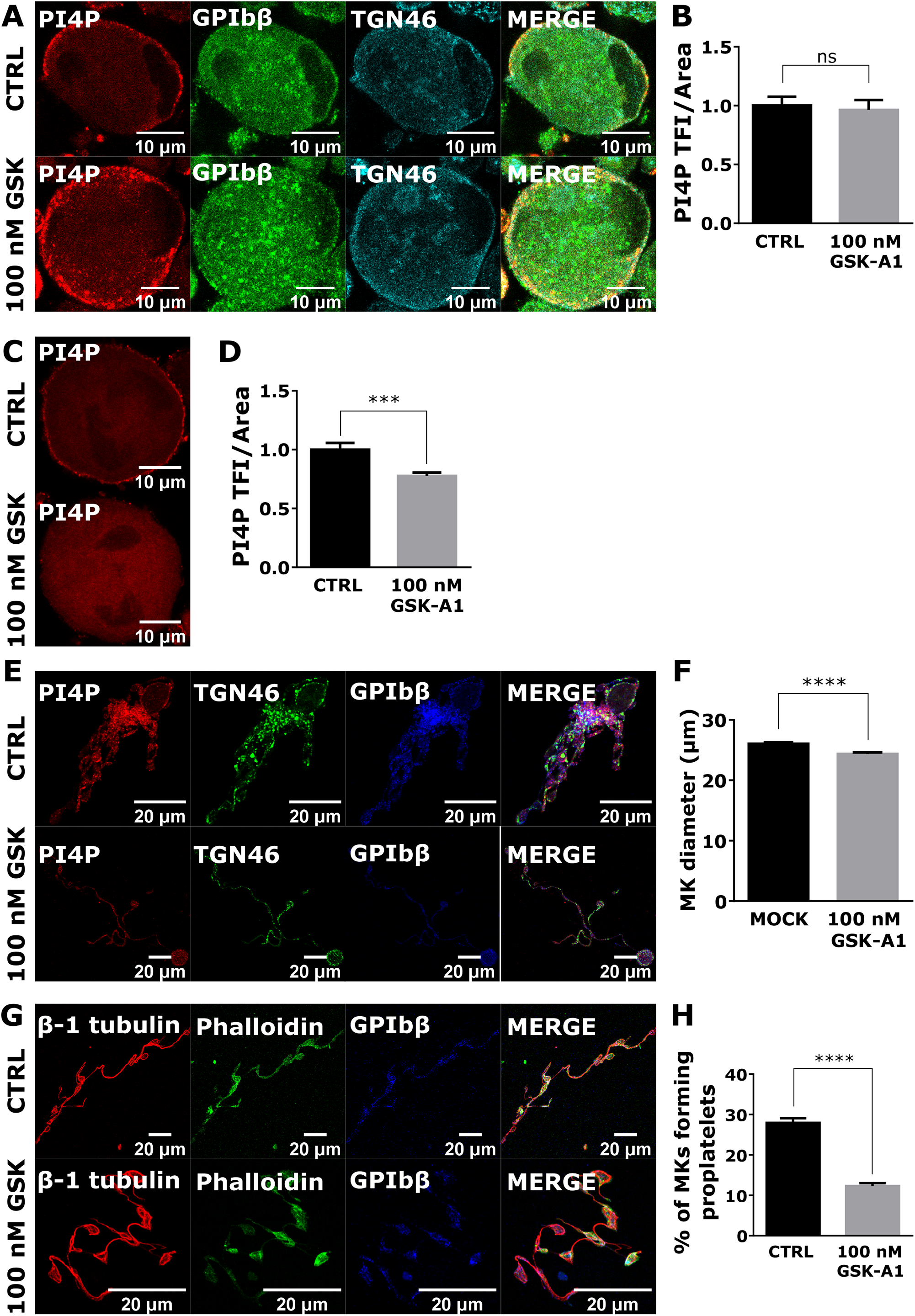
The inhibition of PI4P production at the PM significantly decreases MK ability to form proplatelets. MKs were isolated from the mouse FL. On day 3 of the culture, they were enriched by BSA gradient and treated with 100 nM PI4KIIIα inhibitor GSK-A1. On day 4 of the culture, MKs were fixed, **(A)** stained for the intracellular pool of PI4P and co-stained for GPIbβ and TGN46 or **(C)** stained for the PM pool of PI4P. **(B** and **D)** PI4P total fluorescence intensity was measured using Zen Black software. On day 4 of the culture, **(H)** the percentage of MKs forming proplatelets was counted and proplatelets were fixed, stained for **(E)** PI4P and co-stained for TGN46 or GPIbβ or **(G)** β-1 tubulin and co-stained with phalloidin or GPIbβ. **(F)** The MK diameter was measured using the AxioVision software. The scale bar of the images is **(A** and **C)** 10 μm and **(E** and **G)** 20 μm. Representative images display a single confocal optical section. Results in the graphs are presented as means, error bars denote ± SEM from at least 3 independent experiments. *, p < 0.05; **, p < 0.01; ***, p < 0.001; **** p < 0.0001; n.s., non-significant.

### SACM1L translocates from the Golgi apparatus to the endoplasmic reticulum in a growth factor-dependent manner in the DAMI cell line

To confirm the role of PI4P in MK maturation, we examined SACM1L and PI4P expression and localization in DAMI^19^ cells. We first analyzed if there is any difference in the SACMI1L expression levels in undifferentiated and differentiated DAMI cells during a period from 4 to 48 hours in conditions with and without nutrients. As shown in Figure 5, A and B, we observed two SACM1L isoforms as in primary MKs (67 and 55.5 kDa, consistent with the sizes found in human tissues, https://www.proteinatlas.org/ENSG00000211456-SACM1L). We did not observe any difference in SACM1L expression during DAMI cells cultivation, upon differentiation in growth factors–rich conditions (Figure 5, A and B) or growth factors-poor conditions (Supplemental Figure 3, A and B). Therefore, we concluded that the expression level of SACM1L is stable during DAMI cells cultivation and does not change upon differentiation.

**Figure 5.**
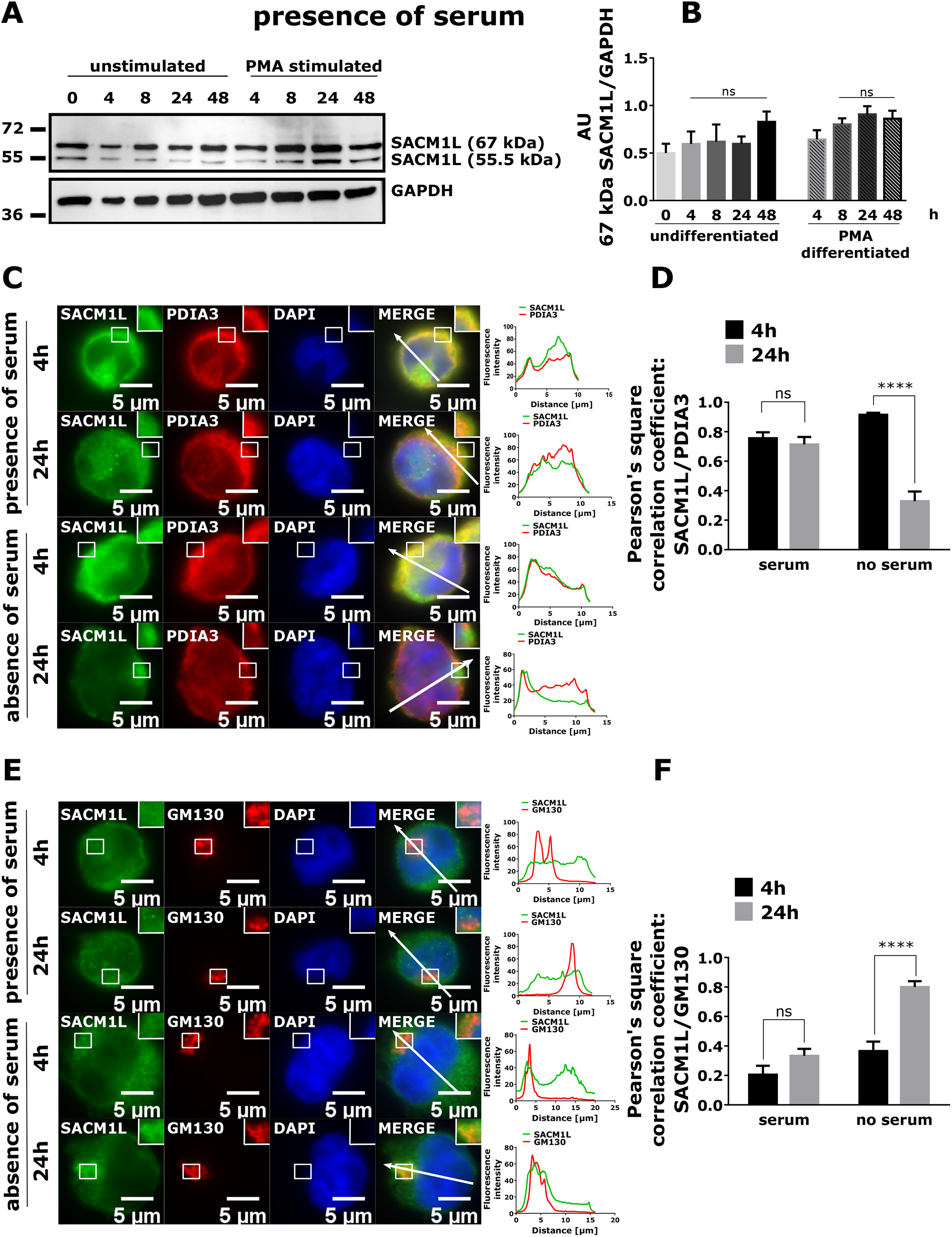
SACM1L shifts from the ER to the Golgi apparatus in a growth factor-dependent manner in undifferentiated DAMI cells. Undifferentiated and PMA (100 nM) differentiated DAMI cells were cultivated **(A** and **B)** in the presence of nutrients for 4, 8, 24, or 48h of nutrients for 4, 8, or 24h. The cells were then lysed, separated by SDS-PAGE, blotted onto a nitrocellulose membrane, and incubated with indicated antibodies. Undifferentiated DAMI cells were cultivated in the presence or absence of nutrients. After 4 or 24 hours they were fixed, stained for SACM1L and co-stained for **(C)** PDIA3 or **(E)** GM130. The scale bar of the images is 5 μm. Cells were analyzed by measuring Pearson’s correlation coefficient through the indicated arrows while the fluorescence intensity graphs show the fluorescence intensity through the same arrows. **(D** and **F)** Results in the graphs are presented as means, error bars denote ± SEM from at least 3 independent experiments. *, p < 0.05; **, p < 0.01; ***, p < 0.001; **** p < 0.0001; n.s., non-significant.

Next, we examined if SACM1L translocates from the Golgi apparatus to the ER in a growth factors-dependent manner. We cultivated undifferentiated DAMI cells for 4 and 24 hours with or without serum, stained them for SACM1L and co-stained them for the marker of the ER (PDIA3) or the Golgi apparatus (GM130). In the presence of serum, SACM1L localizes to the ER (Figure 5C, upper panels) but does not localize to the Golgi apparatus (Figure 5E, upper panels) after 4 and 24 hours of culture. On the other hand, when cells are deprived of growth factors, after 24 SACM1L shifts from the ER (Figure 5C, lower panels) to the Golgi apparatus (Figure 5E, lower panels). The shift is also shown by a decrease in colocalization of SACM1L with the ER (Figure 5D), and an increase in colocalization with the Golgi apparatus (Figure 5F). This suggests that, as shown in fibroblasts^12^, in undifferentiated DAMI cells SACM1L shifts from the ER to the Golgi apparatus in a growth factor-dependent manner and also confirms the specificity of the antibody used.

### Expression of WT and mutant SACM1L in DAMI cell results in a change in PI4P level and localization

We next wanted to study the effect of SACM1L on PI4P by exogenously expressing wild-type and mutant SACM1L in differentiated DAMI cells. Both WT and mutant SACM1L localize to the ER (Supplemental Figure 4), consistent with the localization obtained by staining with the antibody in serum conditions (Figure 5). PI4P is mostly localized at the Golgi in differentiated DAMI cells (Figure 6, top panel). Upon expression of WT SACM1L, the levels of PI4P do not change (Figure 6B) but the localization does change. From its compact localization at the Golgi, PI4P disperses throughout the cell (Figure 6A) in approximately 80% of the cells (Figure 6D). In contrast, the expression of mutant SACM1L significantly increases PI4P levels (Figure 6B) with no change in its localization (Figure 6, A and D). Both WT and C389S SACM1L did not impact the size of DAMI cells (Figure 6C). It is worth mentioning that we observed that undifferentiated DAMI cells were hard to transfect with these constructs so we first stimulated DAMI cells with PMA for 24 hours, and then transfected them with SACM1L constructs which could explain the lack of phenotype on cell size. Since we could not observe a decrease of PI4P with the expression of wild-type SACM1L, we wanted to test if the antibody for PI4P is specific by expressing the P4M-SidM probe^20^ which was shown to specifically bind to PI4P. As shown in Supplemental Figure 5, the P4M-SidM probe localizes at the Golgi where we could observe the staining of PI4P. This suggests that the staining of PI4P is specific and obtained results with no change in PI4P levels could indicate that upon WT SACM1L expression the cells try to compensate for the loss of PI4P by producing it ectopically via different pathways.

**Figure 6.**
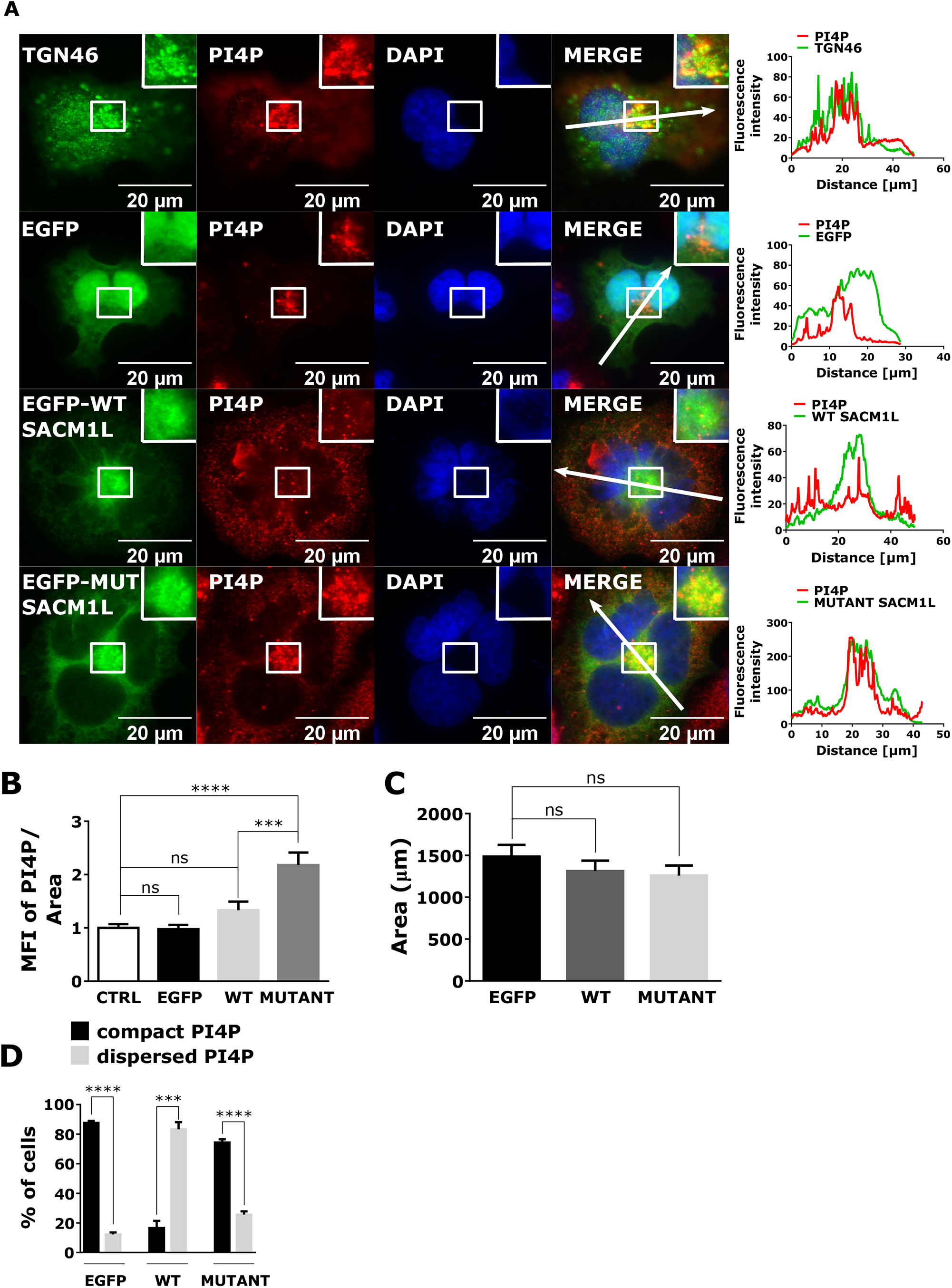
Exogenous expression of wild-type and mutant SACM1L changes PI4P levels and localization in DAMI cells. DAMI cells were differentiated with 100 nM PMA for 24h, transfected for wild-type or mutant SACM1L for an additional 24h, fixed, and stained for PI4P. The cells were analyzed by measuring **(B)** the mean fluorescence intensity of PI4P using FIJI ImageJ software, **(C)** cell area, **(D)** and the percentage of cells that have compact or dispersed PI4P staining. The fluorescence intensity graphs show the fluorescence intensity through the indicated arrows. Results in the graphs are presented as means, error bars denote ± SEM from at least 3 independent experiments. *, p < 0.05; **, p < 0.01; ***, p < 0.001; **** p < 0.0001; n.s., non-significant.

## Discussion

Here we show that PI4P changes location during MK maturation. In young MKs, PI4P is mostly confined to the Golgi apparatus, while in mature MKs it is mostly present at the PM. Both the Golgi and the PM pool of PI4P mediate MK maturation and proplatelet formation since the expression of WT-SACM1L and the inhibition of PI4KIIIα significantly reduce the number of MK forming proplatelets. Although primarily found on the Golgi apparatus, PI4P also resides on the PM and ER in other cells^5^, as well as late endosomes and lysosomes^20^. The role of PI4P at the PM is not clear. It was shown that it can serve as a substrate for the formation of phosphatidylinositol-4,5-bisphosphate [PI(4,5)P_2_], but it can also contribute to the polyanionic lipid pool that defines the inner leaflet of the PM^21^. A nonvesicular lipid transport driven by PI4P concentration gradient was recently described: lipid transfer protein domains at membrane contact sites transport lipid cargo from ER to Golgi or PM, then same transfer domains traffic PI4P from membranes rich in this lipid (Golgi or PM) back to ER where it is being hydrolyzed by SACM1L^22,23^. It is important to note that knock-out of SACM1L in mice is embryonically lethal suggesting a housekeeping role of SACM1L^29^.

In mature MKs in our experiments, it was not expected for PI4P to maintain its perinuclear localization since the Golgi apparatus itself undergoes morphology changes during megakaryopoiesis. In line with our observations, it has been previously shown that immature MKs have a condensed, perinuclear Golgi while in mature MKs the Golgi apparatus is widespread throughout the cell^4,24–27^. The Golgi apparatus may disperse to equip the DMS with needed proteins and lipids. It is possible that the dispersal of Golgi in mature MKs is preceded by the decrease of Golgi PI4P found in immature MKs. Rather, in mature MKs PI4P is present in punctate structures in the periphery and on PM. A similar phenotype with fragmented Golgi and PI4P in punctate structures and peripheral membranes was observed when SACM1L was knocked-down in other cells^28,29^. In our experiments SACM1L is stably expressed at all stages of MK development, however it changes localization: in young MKs, it is notably condensed in the perinuclear region (ER/Golgi) while at later stages SACM1L is scattered throughout the cell and follows ER (Fig. 1). Differential SACM1L localization could lead to intracellular PI4P hydrolysis, however accumulation of PI4P at the PM could be due to PI4KIIIα production. The exogenous expression of WT-SACM1L in MKs results in a retention of the Golgi apparatus perinuclearly and its colocalization with TGN46 which subsequently leads to increased intracellular localization of PI4P (Fig. 3). These results indicate that a balanced amount of functional SACM1L needs to be present in MKs. There is a clear correlation between exogenous SACM1L expression and PI4P/Golgi localization, leading to the formation of smaller MKs that produce significantly fewer proplatelets, while catalytically dead SACM1L had no effect. WT, but not catalytically dead SACM1L, was shown to interact with COPI^30^, therefore excess of WT-SACM1L molecules in MKs could bind additional COPI molecules and favor retrograde transport from Golgi to ER. In addition, an actute PI4P depletion was shown to diminish anterograde trafficking from the Golgi to PM and late endosomes^9^. Both of those scenarios could reduce the dispersion of the Golgi and consequently maturation of WT-SACM1L expressing MK.

The changes in Golgi morphology have been investigated in mice^4,27^, rats^24,25^ or guinea pigs^26^ but not in human MKs. It would be interesting to see if the same applies to human MKs. MKs are not the only cells in which Golgi fragmentation has been observed. Small Golgi fragments have been found in neuronal dendrites that are separated from the perinuclear Golgi and serve as secretory organelles^31^. These neuronal Golgi fragments are called Golgi outposts (GOs)^32^. It has been shown that GOs are important for dendrite growth and branching because GOs ablation reduces dendritic extension and retraction^33^ and they can also interact with cytoskeletal remodelling and motor proteins^31^. Since GOs in neurons are important for dendrite growth, the Golgi fragments in MKs may have a similar role in the growth of the DMS and proplatelets. PI4P is the most important PI for the anterograde Golgi trafficking, and it could serve as a driver of the Golgi fragmentation.

Several PI4P effector proteins have been shown to directly bind to PI4P in mammalian cells^34^. For some of the PI4P effector proteins, it has been shown to be highly expressed in mouse PLTs such as GGA1 (copy number: 1677), GGA3 (copy number: 342), and GOLPH3 (copy number: 1943)^35^. While GGAs are protein adaptors that localize at the TGN and mediate the transport from the Golgi to endosomes^36^, GOLPH3 also localizes to the TGN via binding to PI4P but it is required for the exit of Golgi vesicles and their anterograde trafficking by inducing curvature of the Golgi membranes^37^. It is possible that perturbations in the PI4P levels or localization (e.g. when WT-SACM1L is expressed) disable the Golgi recruitment of GOLPH3 to the TGN which results in a defect of the Golgi anterograde trafficking. What are the exact PI4P effectors that mediate MK maturation remains to be elucidated.

Although DAMI cells exhibit morphological and biochemical characteristics of MKs and PLTs and are used as a model for megakaryopoiesis, they have their flaws. They do not form the DMS, and even though they can form cytoplasmic protrusions and pseudopods, these cannot be compared to true proplatelets^19^. However, they can be used for further analysis on MKs or as a model for the proof of concept. Here, we show that in undifferentiated DAMI cells, SACM1L shifts from the Golgi apparatus to the ER in a growth factors-dependent manner. Although we could not detect differences in overall PI4P levels in primary MKs with either of the SACM1L constructs, the expression of WT-SACM1L in DAMI cells changed PI4P localization while the expression of mutant SACM1L caused accumulation of PI4P, indicating that diverse types of cells could differently regulate PI4P levels. For measuring PI4P levels we used an established immunocytochemistry method^14^ that was validated by expressing a PI4P-binding probe in the DAMI cell line (P4M-SidM). We tried to express different PI4P binding probes (FAPP^8^, OSBP^38^) in MKs, however, the transduction rate was very low (data not shown) which disabled the analysis of PI4P levels and localization with this method.

The lack of difference in PI4P fluorescence intensity levels accompanied by dispersion of PI4P throughout the cell from the compact Golgi localization in WT-SACM1L expressing DAMI (or primary MKs) might be due to ectopic PI4P formation due to low intracellular PI4P levels. It was shown in HeLa cells that the inhibition of PI4KIIIα, a kinase that produces PI4P at the PM, leads to a decrease of PI4P levels at the PM with a simultaneous increase of intracellular PI4P^18^.

In conclusion, we demonstrate that both the intracellular pool of PI4P that is controlled by the SACM1L phosphatase and the PM pool of PI4P that is controlled by the PI4KIIIα kinase are necessary for MKs maturation and proplatelet formation. These results suggest that tight control of the levels and localization of PI4P is important for megakaryopoiesis.

## Acknowledgements

This work was supported by the Croatian Science Foundation (CSF) grant no. UIP-2014-09-2400, ICGEB starting grant no. CRP/HRV15-04_EC, University of Rijeka support 18-188-1343, Croatian Science Foundation support for a PhD student (ESF-HRZZ-01-2018, A.B.), American Society of Hematology Global Research Award (A.J.B.).

## Authorship contributions

A.B. designed and performed the experiments, analyzed the data, and wrote the manuscript; A.J.B. conceived and designed the research, analyzed and interpreted the data, wrote the manuscript, and supervised and handled funding.

## Conflict of interest disclosures

The authors declare no conflict of interest.

## Supplemental Figure legends

**Supplemental Figure 1. The Golgi apparatus disperses during MKs maturation**. MKs were isolated from **(A)** the mouse BM or **(B)** the mouse FL, cultivated for **(A)** 3 and 5 days or **(B)** 2 and 4 days, and enriched by a BSA gradient. The cells were then fixed, stained for GM130 and co-stained for TGN46. Representative images display a single confocal optical section. The scale bar of the images is 10 μm. **(B** and **D)** The number of cells with different Golgi structures was counted. Results in the graphs are presented as means, error bars denote ± SEM from at least 3 independent experiments. *, p < 0.05; **, p < 0.01; ***, p < 0.001; **** p < 0.0001; n.s., non-significant.

**Supplemental Figure 2. FL-derived MKs readily express wild-type and mutant SACM1L**. MKs were isolated from the mouse FL. On day 2 of the culture, they were infected with wild-type or mutant SACM1L. On day 3 of the culture, they were enriched by a BSA gradient. **(A)** On day 4 of the culture, they were imaged with an epifluorescent microscope, **(B)** and the number of MKs expressing different constructs was counted. **(C)** The cells were then fixed and stained for GPIbβ. Representative images display a single confocal optical section. Results in the graphs are presented as means, error bars denote ± SEM from at least 3 independent experiments. *, p < 0.05; **, p < 0.01; ***, p < 0.001; **** p < 0.0001; n.s., non-significant.

**Supplemental Figure 3. SACM1L expression is not growth factors – dependent in DAMI cells**. Undifferentiated and PMA (100 nM) differentiated DAMI cells were cultivated **(A** and **B)** in the absence of nutrients for 4, 8, or 24h. The cells were then lysed, separated by SDS-PAGE, blotted onto a nitrocellulose membrane, and incubated with indicated antibodies. Results in the graphs are presented as means, error bars denote ± SEM from at least 3 independent experiments. *, p < 0.05; **, p < 0.01; ***, p < 0.001; **** p < 0.0001; n.s., non-significant.

**Supplemental Figure 4. Wild-type and mutant SACM1L localize at the ER in PMA differentiated DAMI cells**. DAMI cells were differentiated with 100 nM PMA for 24h and transfected with wild-type and mutant SACM1L for an additional 24h. The cells were then fixed and stained for PDIA3. The scale bar of the images is 20 μm. The fluorescence intensity graphs show the fluorescence intensity through the indicated arrows.

**Supplemental Figure 5. P4M-SidM probe localizes at the TGN**. DAMI cells were differentiated with 100 nM PMA for 24h and transfected with a P4M-SidM PI4P probe for an additional 24h. The cells were then fixed and stained for TGN46. The scale bar of the images is 20 μm. The fluorescence intensity graphs show the fluorescence intensity through the indicated arrows.

## Notes

### Competing Interest Statement

The authors have declared no competing interest.

## References

1 Machlus KR, Italiano JE. The incredible journey: From megakaryocyte development to platelet formation. Journal of Cell Biology 2013; 201: 785–796.

2 Noetzli LJ, French SL, Machlus KR. New Insights Into the Differentiation of Megakaryocytes From Hematopoietic Progenitors. ATVB 2019; 39: 1288–1300.

3 Boscher J, Guinard I, Eckly A, Lanza F, Léon C. Blood platelet formation at a glance. J Cell Sci 2020; 133. doi:10.1242/jcs.244731.

4 Eckly A, Heijnen H, Pertuy F, Geerts W, Proamer F, Rinckel J-Y et al. Biogenesis of the demarcation membrane system (DMS) in megakaryocytes. Blood 2014; 123: 921–930.

5 Balla T. Phosphoinositides: Tiny Lipids With Giant Impact on Cell Regulation. Physiological Reviews 2013; 93: 1019–1137.

6 Wang YJ, Wang J, Sun HQ, Martinez M, Sun YX, Macia E et al. Phosphatidylinositol 4 phosphate regulates targeting of clathrin adaptor AP-1 complexes to the Golgi. Cell 2003; 114: 299–310.

7 Wang J, Sun H-Q, Macia E, Kirchhausen T, Watson H, Bonifacino JS et al. PI4P promotes the recruitment of the GGA adaptor proteins to the trans-Golgi network and regulates their recognition of the ubiquitin sorting signal. Molecular biology of the cell 2007; 18: 2646–2655.

8 Godi A, Campli AD, Konstantakopoulos A, Tullio GD, Alessi DR, Kular GS et al. FAPPs control Golgi-to-cell-surface membrane traffic by binding to ARF and PtdIns(4)P. Nature Cell Biology 2004; 6: 393–404.

9 Szentpetery Z, Varnai P, Balla T. Acute manipulation of Golgi phosphoinositides to assess their importance in cellular trafficking and signaling. Proceedings of the National Academy of Sciences 2010; 107: 8225–8230.

10 Hsu F, Mao Y. The structure of phosphoinositide phosphatases: Insights into substrate specificity and catalysis. Biochimica et Biophysica Acta (BBA) - Molecular and Cell Biology of Lipids 2015; 1851: 698–710.

11 Del Bel LM, Brill JA. Sac1, a lipid phosphatase at the interface of vesicular and nonvesicular transport. Traffic 2018; 19: 301–318.

12 Blagoveshchenskaya A, Cheong FY, Rohde HM, Glover G, Knödler A, Nicolson T et al. Integration of Golgi trafficking and growth factor signaling by the lipid phosphatase SAC1. Journal of Cell Biology 2008; 180: 803–812.

13 Burkhart JM, Vaudel M, Gambaryan S, Radau S, Walter U, Martens L et al. The first comprehensive and quantitative analysis of human platelet protein composition allows the comparative analysis of structural and functional pathways. Blood 2012; 120: 73–82.

14 Hammond GRV, Schiavo G, Irvine RF. Immunocytochemical techniques reveal multiple, distinct cellular pools of PtdIns4P and PtdIns(4,5)P2. Biochemical Journal 2009; 422: 23–35.

15 Bura A, Begonja AJ. Imaging of Intracellular and Plasma Membrane Pools of PI(4,5)P2 and PI4P in Human Platelets. Life 2021; 11: 20.

16 Schindelin J, Arganda-Carreras I, Frise E, Kaynig V, Longair M, Pietzsch T et al. Fiji: an open-source platform for biological-image analysis. Nat Methods 2012; 9: 676–682.

17 Bertović I, Bura A, Jurak Begonja A. Developmental differences of in vitro cultured murine bone marrow- and fetal liver-derived megakaryocytes. Platelets 2021; : 1–13.

18 Nakatsu F, Baskin JM, Chung J, Tanner LB, Shui G, Lee SY et al. PtdIns4P synthesis by PI4KIIIα at the plasma membrane and its impact on plasma membrane identity. Journal of cell biology 2012; 199: 1003–1016.

19 Sheryl M, Greenberg DS, Rosenthal TA, Greeley RT, Handin RI. Characterization of a New Megakaryocytic Cell Line: The Dami Cell. Blood 1988; 72: 1968–1977.

20 Hammond GRV, Machner MP, Balla T. A novel probe for phosphatidylinositol 4-phosphate reveals multiple pools beyond the Golgi. Journal of Cell Biology 2014; 205: 113–126.

21 Hammond GRV, Fischer MJ, Anderson KE, Holdich J, Koteci A, Balla T et al. PI4P and PI(4,5)P2 Are Essential But Independent Lipid Determinants of Membrane Identity. Science 2012; 337: 727–730.

22 de Saint-Jean M, Delfosse V, Douguet D, Chicanne G, Payrastre B, Bourguet W et al. Osh4p exchanges sterols for phosphatidylinositol 4-phosphate between lipid bilayers. Journal of Cell Biology 2011; 195: 965–978.

23 Zewe JP, Wills RC, Sangappa S, Goulden BD, Hammond GR. SAC1 degrades its lipid substrate PtdIns4P in the endoplasmic reticulum to maintain a steep chemical gradient with donor membranes. eLife 2018; 7: e35588.

24 Behnke O. An electron microscope study of the megacaryocyte of the rat bone marrow. Journal of Ultrastructure Research 1968; 24: 412–433.

25 MacPherson GG. Development of megakaryocytes in bone marrow of the rat: an analysis by electron microscopy and high resolution autoradiography. Proceedings of the Royal Society of London 1971; 177: 265–274.

26 Paulus J-M. DNA Metabolism and Development of Organelles in Guinea-Pig Megakaryocytes: A Combined Ultrastructural, Autoradiographic and Cytophotometric Study. Blood 1970; 35: 298–311.

27 Wandall HH, Rumjantseva V, Sørensen ALT, Patel-Hett S, Josefsson EC, Bennett EP et al. The origin and function of platelet glycosyltransferases. Blood 2012; 120: 626–635.

28 Cheong FY, Sharma V, Blagoveshchenskaya A, Oorschot VMJ, Brankatschk B, Klumperman J et al. Spatial Regulation of Golgi Phosphatidylinositol-4-Phosphate is Required for Enzyme Localization and Glycosylation Fidelity. Traffic 2010; 11: 1180–1190.

29 Liu Y, Boukhelifa M, Tribble E, Morin-Kensicki E, Uetrecht A, Bear JE et al. The Sac1 Phosphoinositide Phosphatase Regulates Golgi Membrane Morphology and Mitotic Spindle Organization in Mammals. MBoC 2008; 19: 3080–3096.

30 Rohde HM, Cheong FY, Konrad G, Paiha K, Mayinger P, Boehmelt G. The Human Phosphatidylinositol Phosphatase SAC1 Interacts with the Coatomer I Complex. Journal of Biological Chemistry 2003; 278: 52689–52699.

31 Wang J, Fourriere L, Gleeson PA. Local Secretory Trafficking Pathways in Neurons and the Role of Dendritic Golgi Outposts in Different Cell Models. Front Mol Neurosci 2020; 13: 597391.

32 Horton AC, Ehlers MD. Dual Modes of Endoplasmic Reticulum-to-Golgi Transport in Dendrites Revealed by Live-Cell Imaging. J Neurosci 2003; 23: 6188–6199.

33 Ye B, Zhang Y, Song W, Younger SH, Jan LY, Jan YN. Growing Dendrites and Axons Differ in Their Reliance on the Secretory Pathway. Cell 2007; 130: 717–729.

34 Santiago-Tirado FH, Bretscher A. Membrane-trafficking sorting hubs: cooperation between PI4P and small GTPases at the trans-Golgi network. Trends in Cell Biology 2011; 21: 515–525.

35 Zeiler M, Moser M, Mann M. Copy Number Analysis of the Murine Platelet Proteome Spanning the Complete Abundance Range. Molecular & Cellular Proteomics 2014; 13: 3435–3445.

36 Bonifacino JS. The GGA proteins: adaptors on the move. Nat Rev Mol Cell Biol 2004; 5: 23–32.

37 Rahajeng J, Kuna S. R, Makowski L. S, Tran T. T. T, Buschman D. M, Li S et al. Efficient Golgi Forward Trafficking Requires GOLPH3-Driven, PI4P-Dependent Membrane Curvature. Developmental Cell 2019; 50: 573–585.

38 Mesmin B, Bigay J, Moser von Filseck J, Lacas-Gervais S, Drin G, Antonny B. A Four-Step Cycle Driven by PI(4)P Hydrolysis Directs Sterol/PI(4)P Exchange by the ER-Golgi Tether OSBP. Cell 2013; 155: 830–843.

