## Supplemental figures for "The intracellular and plasma membrane pools of PI4P control megakaryocyte maturation and proplatelet formation"

### Supplemental Figure 1

**A**

**BMMK**

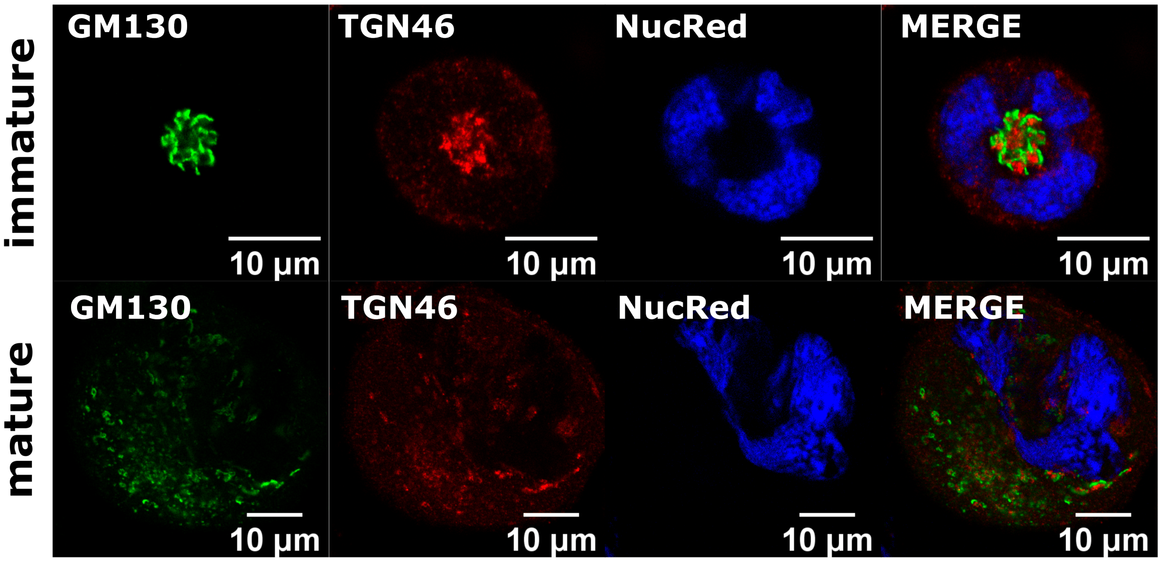


**B**

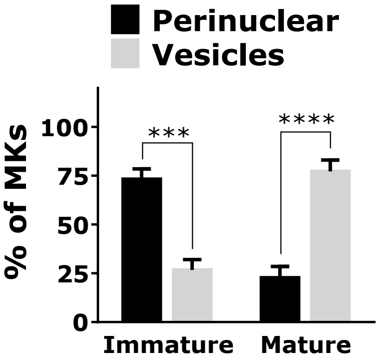


**C**

**FLMK**

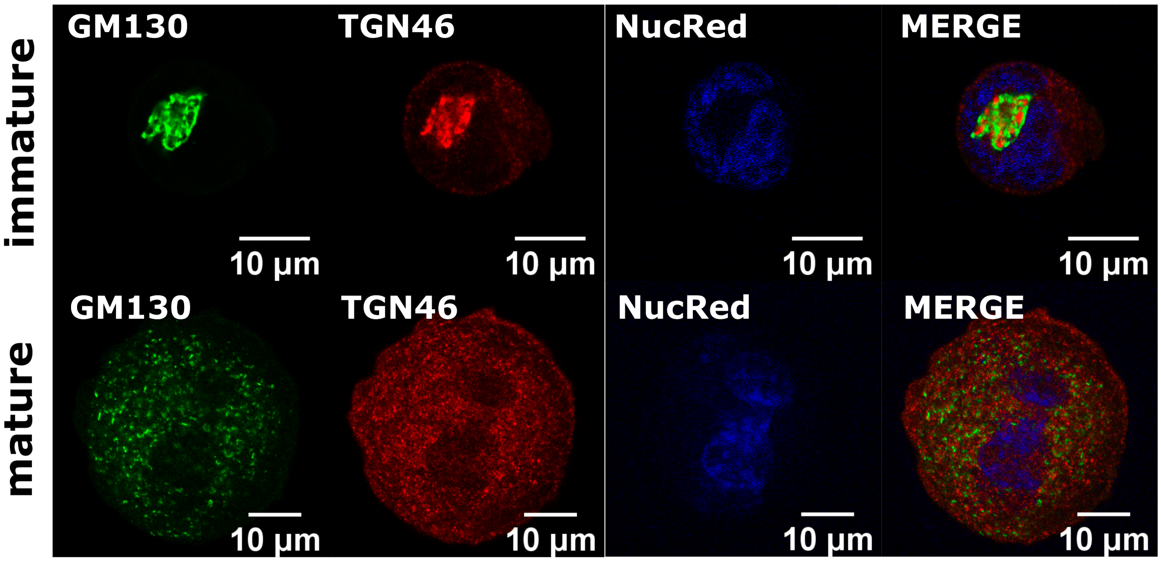


**D**

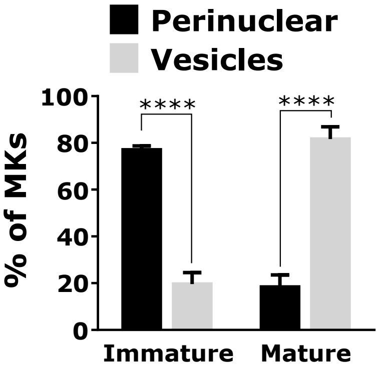

#### Supplemental Figure 2

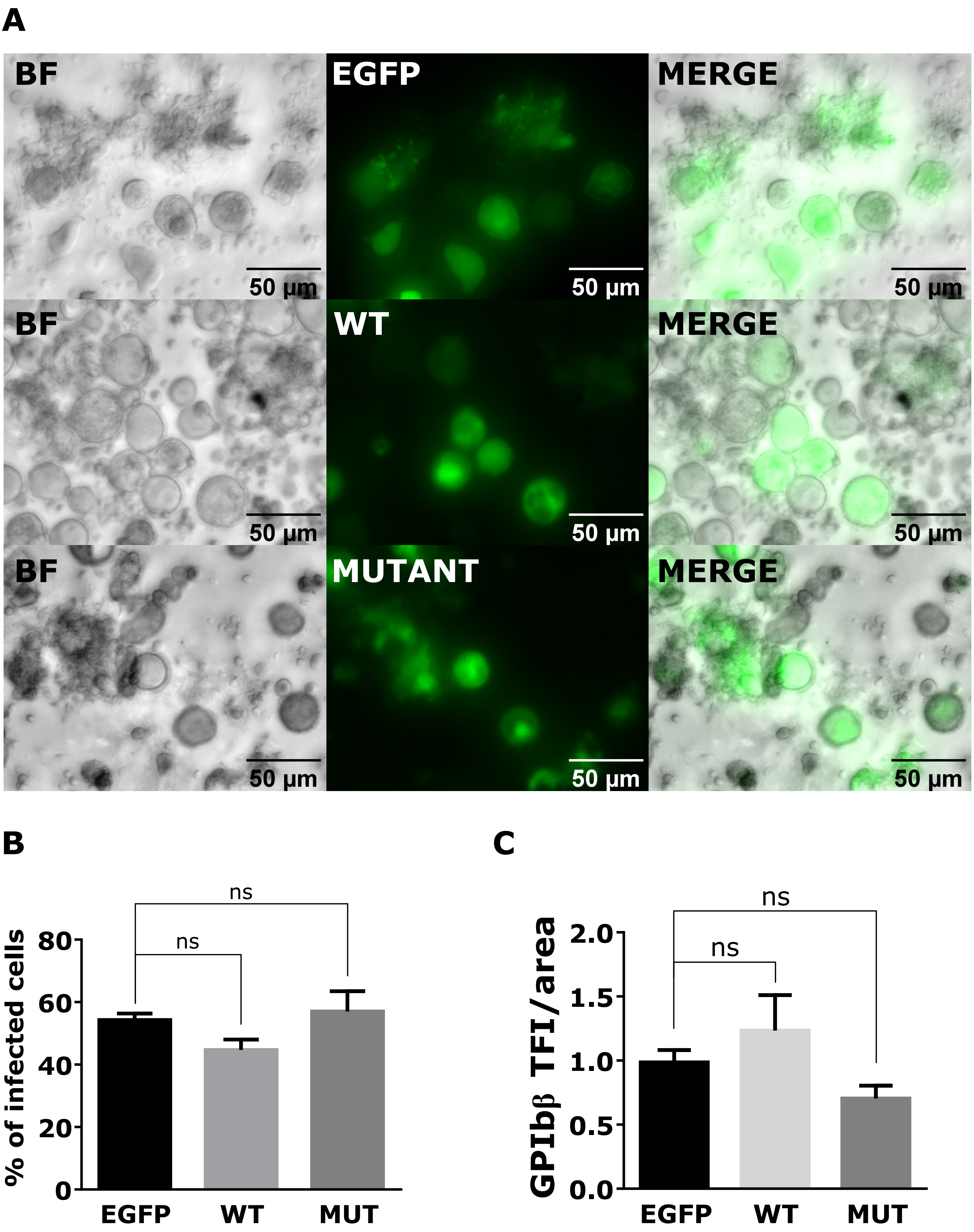

### Supplemental Figure 3

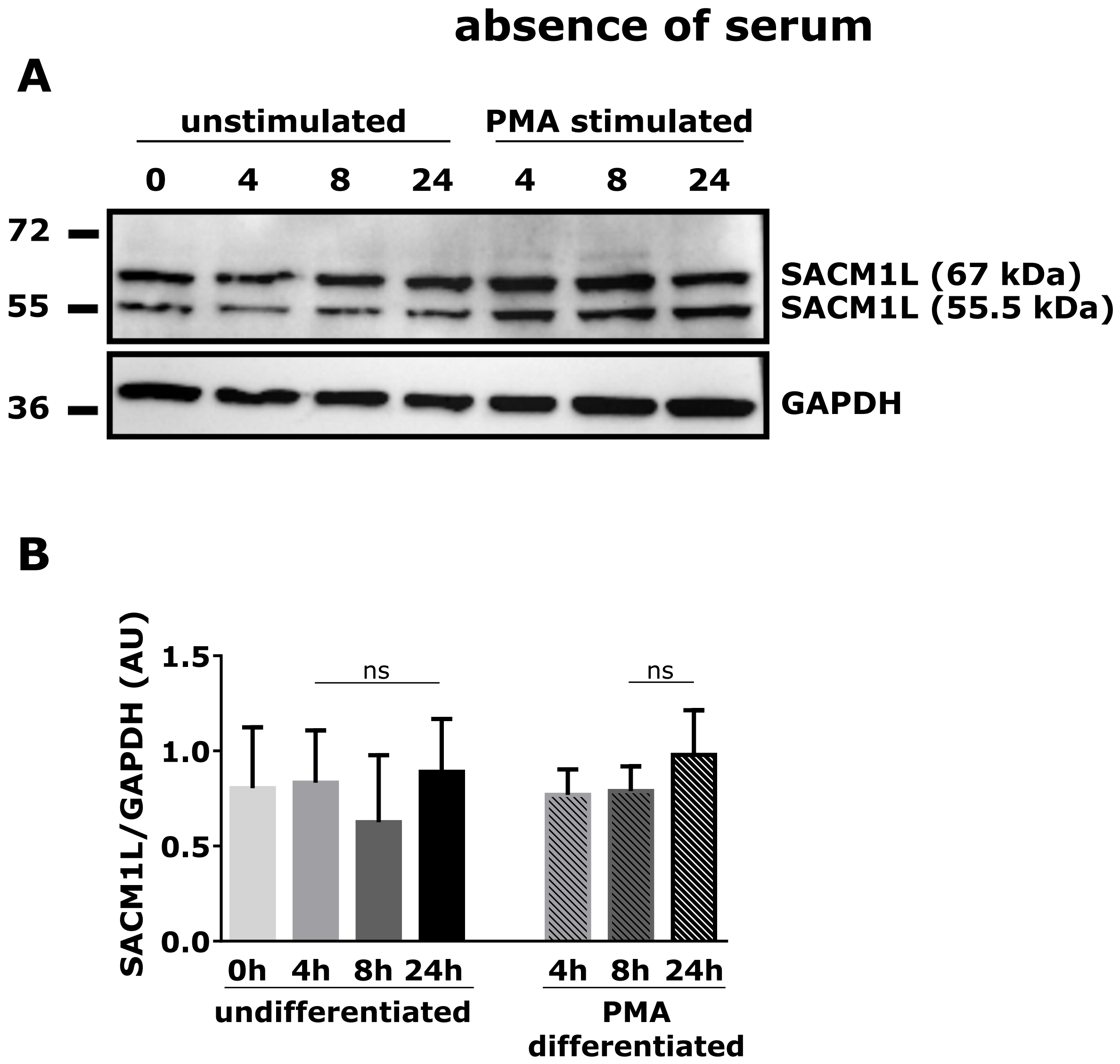

### Supplemental Figure 4

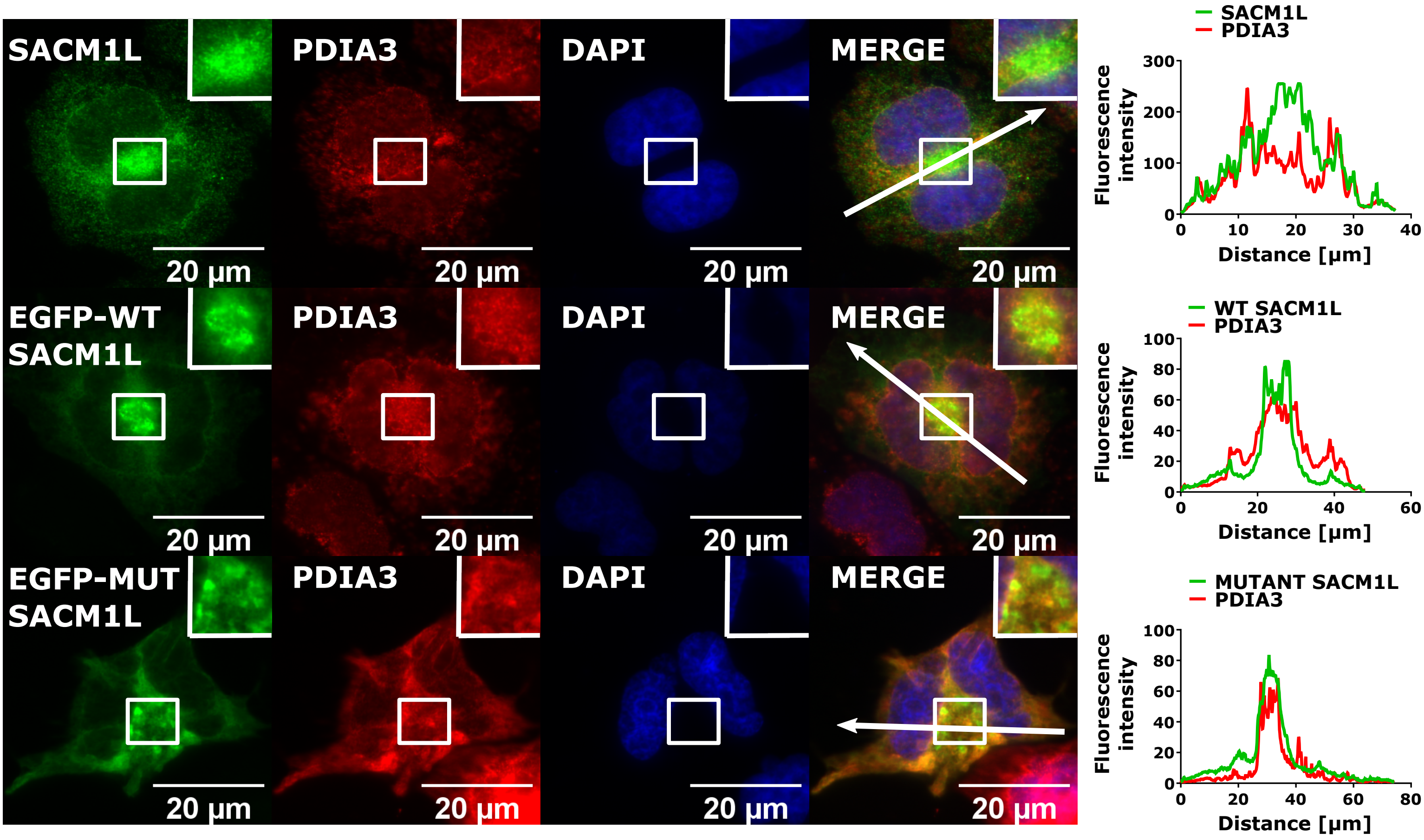

### Supplemental Figure 5

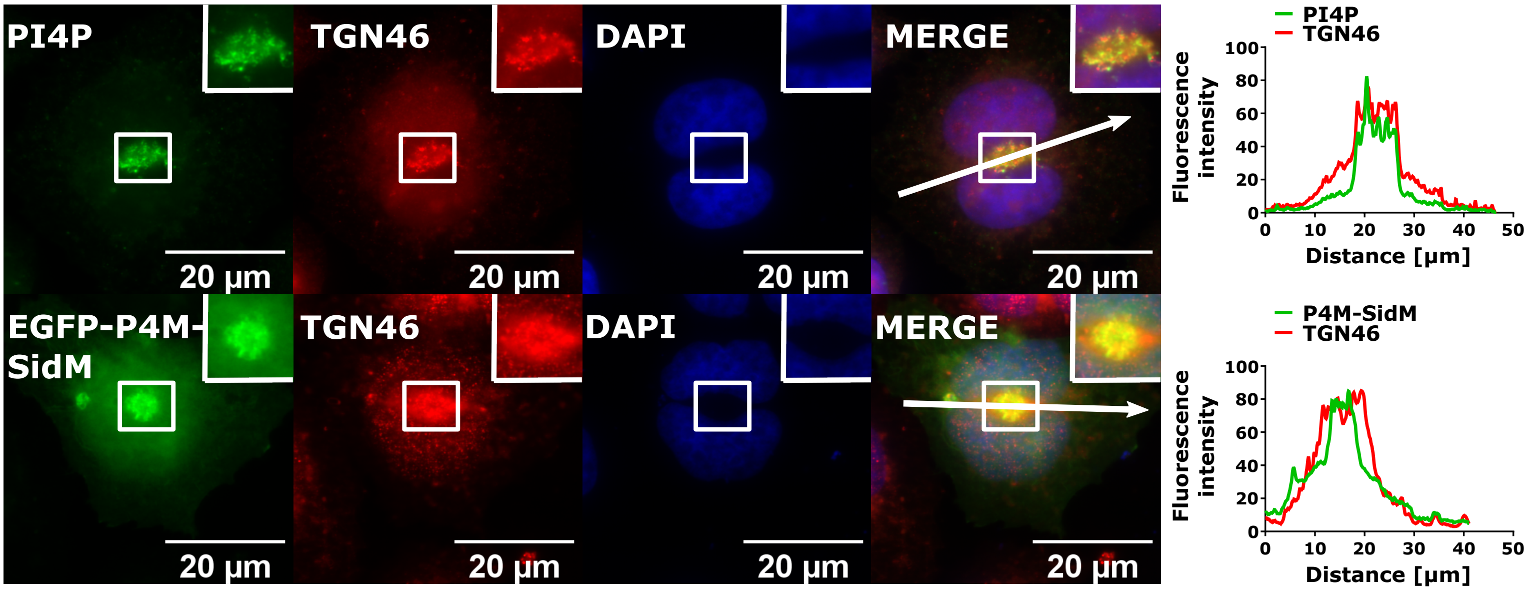
